# Sharing of weak signals of positive selection across the genome

**DOI:** 10.1101/2020.04.22.055954

**Authors:** Nathan S. Harris, Alan R. Rogers

## Abstract

Selection in humans often leaves subtle signatures at individual loci. Few studies have measured the extent to which these signals are shared among human populations. Here a new method is developed to compare weak signals of selection in aggregate across the genome using the 1000 Genomes Phase 3 Data. Results presented here show that selection producing weak selection serves to increase population differences around coding areas of the genome.

## 1 Introduction and background

Until relatively recently, studies of natural selection in humans focused on classic selective sweeps that have large effects on isolated regions of the genome (Sabeti et al., 2002; Voight et al., 2006). In a classic selective sweep, a new beneficial mutation appears in one person and spreads through a population (Smith and Haigh, 1974). When an allele is sweeping through the population, surrounding DNA from the original haplotype on which the mutation occurred tends to “hitchhike” with the selected allele. This results in linkage disequilibrium (LD), a nonrandom association of alleles at two or more loci (Lewontin and Kojima, 1960). The blocks around selected loci are longer and contain less diversity the greater the strength of selection. Mutation and recombination reintroduce variation into blocks of LD (Lande, 1975), and given sufficient generations following the sweep, LD blocks around a locus are broken apart. The extent to which this has occurred depends on the local mutation and recombination rates and the amount of time that has passed. This “signal” is used by a variety of methods to detect natural selection (Booker et al., 2017; Haasl and Payseur, 2016; Vitti et al., 2013).

As research continues it has become apparent that the most common forms of selection in humans are those that have smaller effects on LD around individual loci such as polygenic adaptation (Daub et al., 2013; Hernandez et al., 2011; Pritchard et al., 2010) and selection on existing variation (Harris et al., 2018; Schrider and Kern, 2017). These forms of selection are unlikely to generate significant signals using statistics designed for classic sweeps, although they may account for some fraction of those that fail to reach significance. Furthermore, while classic selective sweeps are usually geographically local and therefore tend to increase population differences (Fagny et al., 2014; Vitti et al., 2013), much less is known about how often weak signals of selection are shared between populations. Weak signals of selection are more likely to be shared between populations because the types of weak selection that produce them are slower, and alleles that arose in a common ancestor are more likely to still be polymorphic. In the context of this paper, a “weak” signal refers to signals of selection that do not reach significance or an Integrated Haplotype Score (|iHS|) greater than two. This term is therefore relative to the statistic being calculated rather than reflecting a rigid category of selective pressure.

Theoretically, populations ought to share more signals of selection if they are closely related for the following reasons:

1. Beneficial mutations are more likely to become lost or fixed the more time that has elapsed since mutation. If two populations are distantly related, beneficial mutations are likely to be either fixed or lost in one or both of the populations. If two populations are closely related, signals from completed sweeps in their common ancestor are more likely to be preserved and detectable in each. Furthermore, the same beneficial mutation may still be polymorphic and increasing in frequency in both populations, producing a shared signal.
2. Closely related populations often share similar environments. If each population experiences a beneficial mutation near the same locus, both populations may show evidence of selection in the same region.
3. Neutral mechanisms can produce spurious signals of selection by chance. Closely related populations are expected to share such signals because a larger portion of their population history is shared.

This theoretical expectation has been supported empirically. Pickrell et al. (2009) show that populations within the same continent are more likely to share signals of selection. Similarly, Johnson and Voight (2018) found that regions of the genome with high concentrations of large |iHS| scores were more likely to overlap between populations if those populations are closely related.

Here, methods traditionally used to detect classic selective sweeps are implemented to characterize genome-wide patterns of shared weak signals of selection. While methods have been developed to detect subtler signatures of selection (Field et al., 2016; Schrider and Kern, 2016), some of the methods for detecting classic selective sweeps still have some utility for studying population differences. Hard selective sweeps may be an uncommon mechanism of adaptation in humans (Coop et al., 2009; Hernandez et al., 2011; Schrider and Kern, 2017), but they are an important case. Many large adaptive changes have occurred in the recent past via classic selective sweeps (Fagny et al., 2014; Vitti et al., 2013). These signals are known to be recent because ancient adaptation that occurred via this mechanism has already driven the causative variant to fixation and the signal has been obscured by subsequent recombination and mutation. It may be possible to detect older instances of selection with some of the same methods because moderately beneficial alleles increase in frequency much slower and the resulting signal persists longer. However, because |iHS| is standardized, and most large significant signals are the result of selection within the last 20,000 years (Voight et al., 2006), ancient signals are unlikely to produce significant signals of selection at individual loci. For this reason, this research investigates genome-wide patterns of weak signals of selection rather than identifying particular loci under selection.

## 2 Results

### 2.1 Weak signals of selection

To get an idea of the relevant strength of selection, a catalog of significant |iHS| signals for the 1000 Genomes Phase 3 data were obtained from Johnson and Voight (2018). These signals were binned by their size to search for any clear cutoffs in the size (in base pairs) of selection signals from |iHS| (Figure 1). Most significant |iHS| signals were at least 100kb long. A model was adapted from Gillespie (2004) to determine the relationship between the selection coefficient (s), the recombination rate (r), and the size of linkage disequilibrium blocks around a selected allele at an intermediate frequency of 0.5 (Figure 2). The strongest signals of classic sweeps will therefore commonly have a selection coefficient of 0.01 or greater. Here we hoped to exclude the majority of these signals by removing sites with |iHS| values greater than two. The remaining sites should disproportionately be from loci with selection coefficients smaller than 0.01.

**Figure 1:**
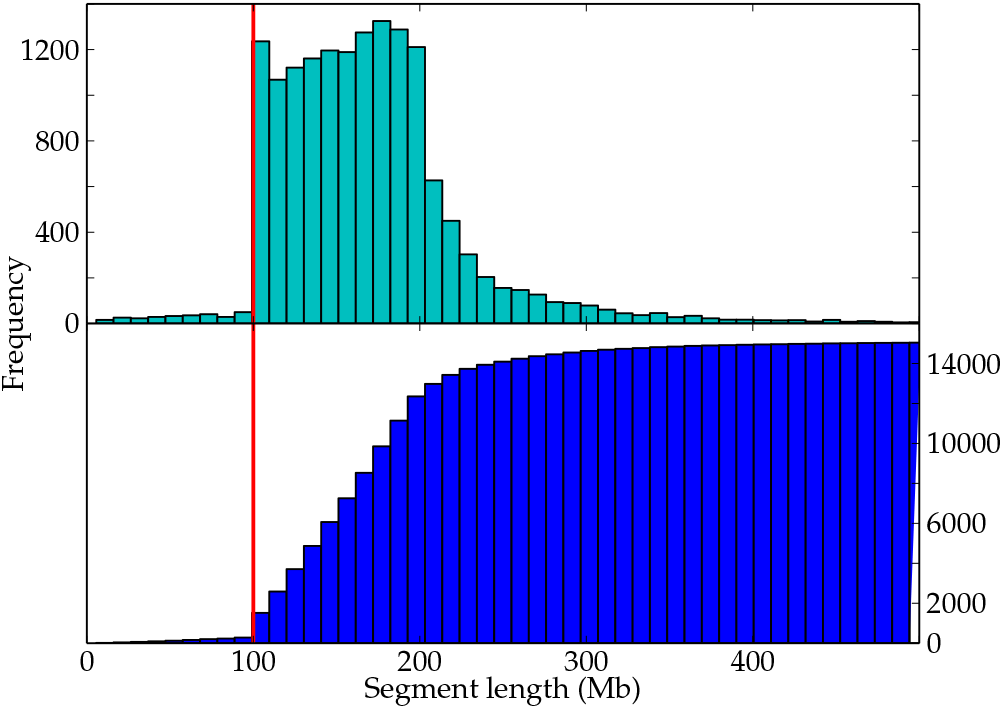
Normal and cumulative histogram of significant |iHS| region sizes in 26 1000 Genomes Phase 3 data. A sharp cliff occurs at 100kb, implying most cases of classic selective sweeps considered to be significant leave signals grearelevantter than 100kb.

**Figure 2:**
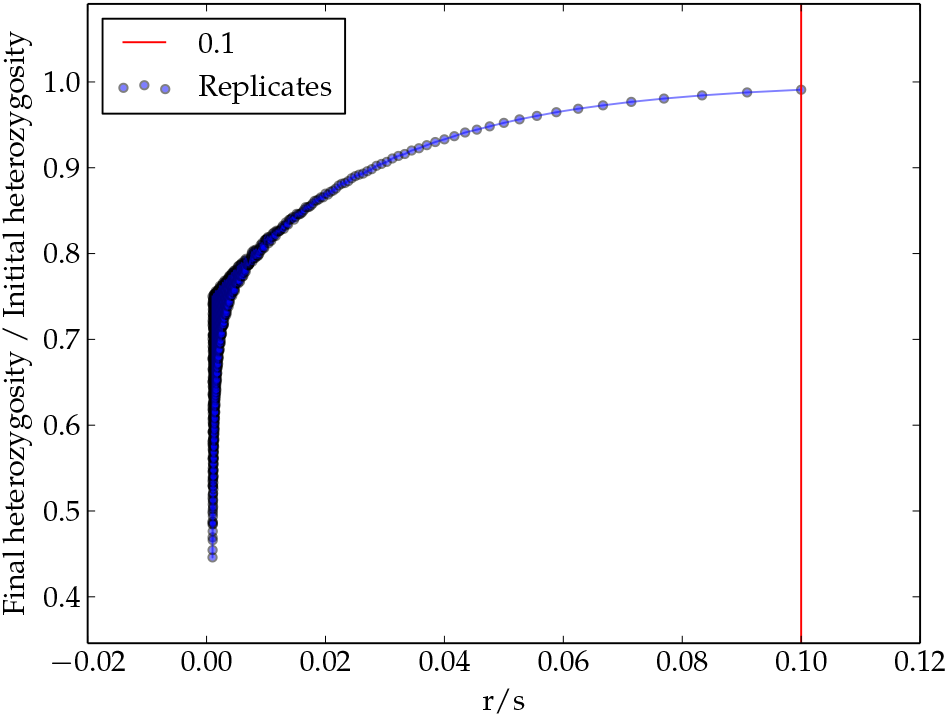
Determining the relationship between the strength of selection and LD block size. The recombination rate (r) is held constant while the selection coefficient (s) varies between replicates. Each replicate is allowed to run until the beneficial allele reaches a frequency of 0.5. The red line indicates the point at which sites are no longer in LD with the site under selection. With known r and block size, s can be determined.

### 2.2 The Integrated Haplotype Score (|iHS|)

To measure signals of selection, |iHS| was calculated for samples from the 1000 Genomes Project. |iHS| identifies regions under selection by comparing the difference in LD between carriers of the reference and alternate alleles. Large |iHS| scores indicate a substantial difference in LD. Correlation of sample |iHS| scores of two populations was calculated for each pair of samples. This correlation was calculated separately for genic and nongenic portions of the genome. Unlike genetic drift, selection affects specific loci and linked variation rather than the entire genome. Genic regions should more commonly be the target of selection because they are more often functional (Barreiro et al., 2008; Coop et al., 2009). The difference between genic and nongenic correlations at a given value of genetic distance is likely the result of selection. While selection is known to occur in some noncoding regions (Forni et al., 2014; Hernandez et al., 2011; Ponjavic et al., 2007), this should have the effect of making any observed differences between genic and nongenic regions more conservative. In either case, correlation is expected to be relatively large and positive when the two populations have both experienced selection in the same areas of the genome. This occurs not only because particular SNPs may be under selection, but also because |iHS| scores at linked sites near mutually selected loci should covary as well.

To limit the effect of hard sweeps, loci with significant |iHS| values (greater than two) were removed from the calculation. To compensate for the effects of linkage, regions were considered nongenic if they were at least 500kb from genic regions. The resulting correlations were regressed with a Loess algorithm against Nei’s genetic distance for each pair of populations. Confidence intervals were generated using a moving block bootstrap (Liu and Singh, 1992) with a block size of 500kb. The outcome is shown in Figure 3.

**Figure 3:**
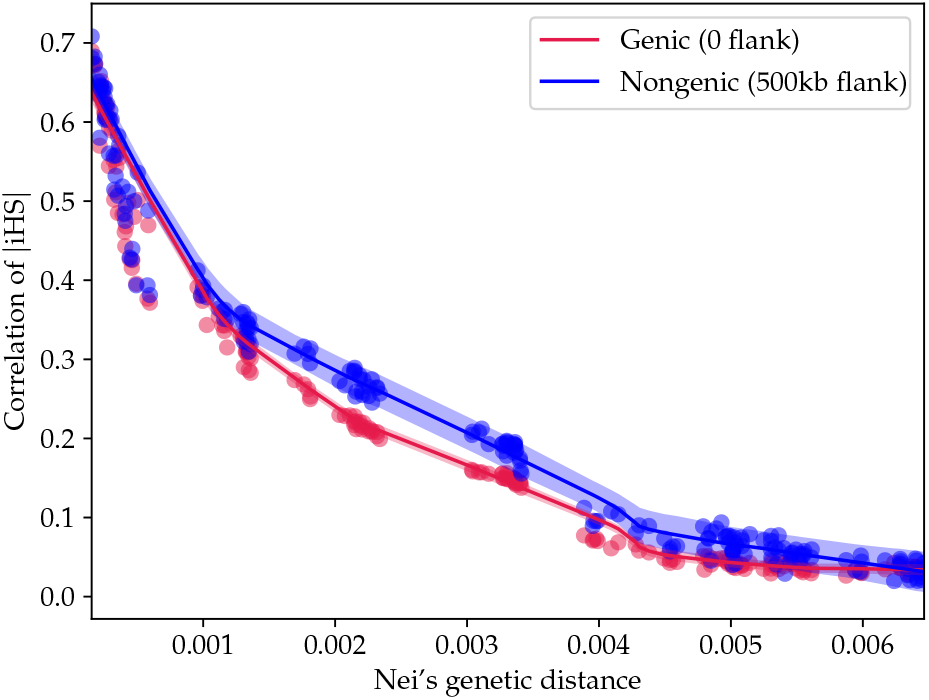
Correlation of |iHS| scores for nongenic regions compared to genic regions.

The left edge of the graph refers to pairs of populations that are genetically similar and tend also to be geographically close. Samples from the same geographic regions experience similar correlations in genic and nongenic sections of their genome. As the genetic distance between samples increases, the correlation between samples in both genic and nongenic regions decreases until it approaches zero in the most genetically distant comparisons, Africans and Eastern Asians. However, the correlation of |iHS| in genic and nongenic regions does not decrease at the same rate. The pairwise genic correlation is consistently lower than the pairwise nongenic correlation for all sample pairs except at large genetic distances.

### 2.3 The Cross Population Extended Haplotype Homozygosity (XP-EHH)

The above analysis was repeated for XP-EHH. XP-EHH compares LD differences between two samples, identifying where selection has occurred in one sample but not the other. XP-EHH was chosen as a supplementary analysis to |iHS| because it is more sensitive to selection in which the beneficial is near fixation. Like |iHS|, XP-EHH correlation between populations decreases with increasing genetic distance. Unlike iHS, XP-EHH genic and nongenic correlations overlap considerably (see Figure 4). However, XP-EHH correlations in either nongenic or genic regions were consistently larger than their |iHS| counterparts.

**Figure 4:**
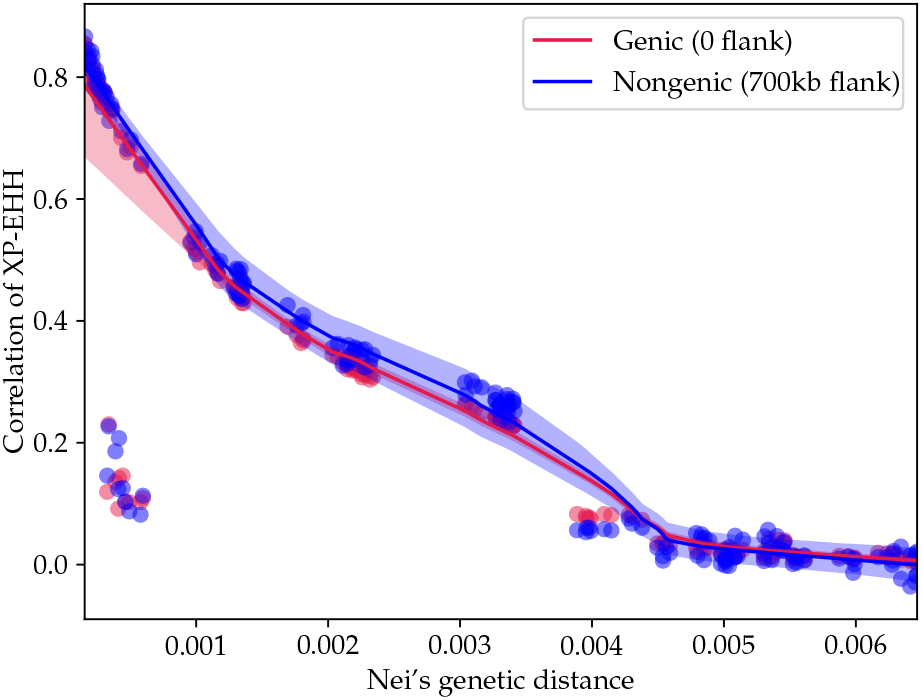
Correlation of XP-EHH scores for nongenic regions compared to genic regions.

### 2.4 Simulations in SLiM

Correlations tend to be lower in genic than in nongenic regions. I will argue that this implies that, even when selection is weak, its primary effect is disruptive, tending to increase differences between populations. To establish this point, simulations were conducted using Selection on Linked Mutations (SLiM) to show that disruptive selection tends to reduce the correlation between |iHS| signals. In each simulation, a single population splits into two. The time of this split varies among simulations to model differences in genetic distance. Following the split, populations experience one of three evolutionary scenarios: neutrality, positive selection, or purifying selection. All three models experience neutral mutations. In both cases of selection, non-neutral mutations occur in a single population following the split. Results were standardized against the neutral simulations with the same divergence time, and the correlation analysis proceeds in the same manner as the real data.

The results of this process are shown in Figure 5 and 6. For any particular divergence time, confidence intervals of neutral and selection models overlapped. However, there is a difference between the scenarios when points are considered together. In each case a Wilcoxon signed-rank test was conducted to assess if the selection correlation could have been drawn from the same distribution as the neutral correlation. The correlation of |iHS| in the presence of positive selection was significantly lower than in the neutral model (*n*=21, *p*=0.002). Previous work has suggested that |iHS| is not sensitive to purifying selection (Enard et al., 2014) and that conclusion was supported here. The correlation of |iHS| in the presence of purifying selection was indistinguishable from the neutral model (*n*=21, *p*=0.357).

**Figure 5:**
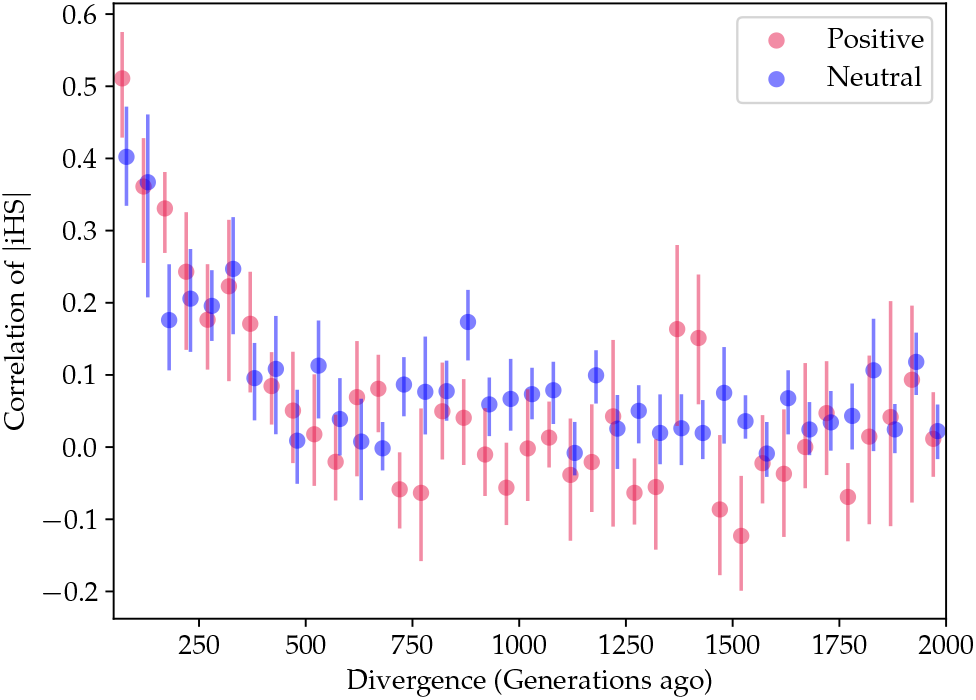
|iHS| correlation with positive selection occurring in one branch. Wilcoxon signed-rank test, *n*=21, *p*=0.002.

**Figure 6:**
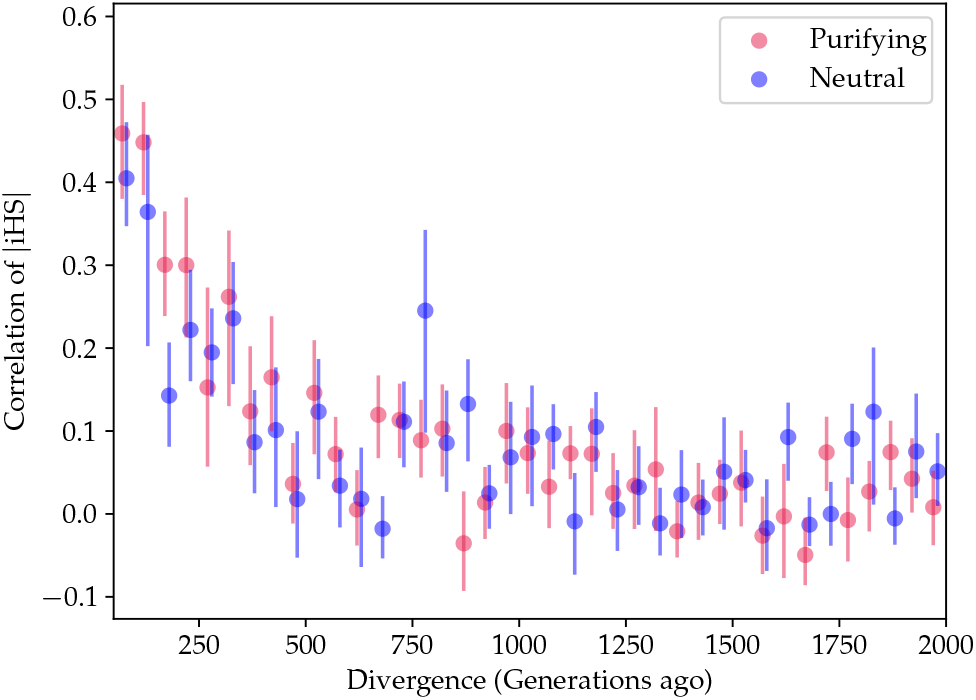
|iHS| correlation with purifying selection occurring in one branch. Wilcoxon signed-rank test, *n*=21, *p*=0.357.

To get an idea of how far back selection can be detected, the basic model of positive selection was repeated for increasing divergence times. In these simulations, beneficial mutations (s=0.01) are introduced into one population following the population split for 2,000 generations. At this point, no more beneficial mutations are introduced and selection on existing beneficial mutations is halted, allowing any existing signals to decay. When divergence times range from 2,000 to 4,000 generations ago (about 50-100-kya) (Figure 7), correlations are small relative to the recent divergence times discussed above, but correlations in selected regions are still significantly smaller than in neutral regions (*n*=40, *p*=2.07e-4). If divergence times are increased further (Figure 8), the significant difference between selected and neutral regions disappears (*n*=40, p=0.83). This strongly suggests that the correlation of |iHS| is sensitive to instances of selection that are much older than the effective range of traditional use of |iHS| (Voight et al., 2006).

**Figure 7:**
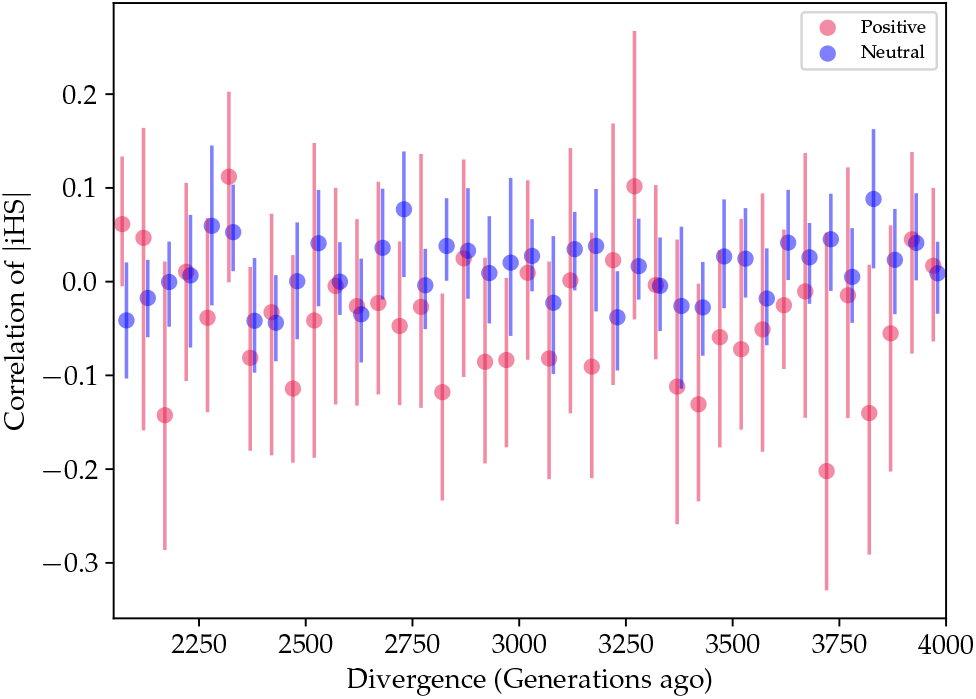
Correlation of |iHS| for a set of divergence times ranging from 2,000 to 4,000 generations ago. While neutral regions have a correlation around zero, selection has the effect of decreasing correlation further.

Simulations of positive selection were repeated using a model of soft selective sweeps, in which a mutation becomes beneficial after it has already drifted to a relatively high frequency. Soft sweeps introduce initial allele frequency as an additional dimension to the simulations. While |iHS| is sensitive to soft sweeps with starting allele frequencies as large as 0.1 (Ferrer-Admetlla et al., 2014), it is unclear to what extent soft selective sweeps will be detected by the methods proposed above. To determine the relevant range of soft sweep parameters, simulations were run over a wide range of combinations of starting allele frequencies and sweep frequency. Soft sweeps, in general, appear to have the same effect as classic sweeps from de novo mutations, depressing the correlation of |iHS| in regions that have experienced selection. The largest allele frequency for which soft sweeps caused a significant difference between selected and neutral regions in our simulations was the 0.09-0.1 bin (*n*=40, p=0.041). The effect size is small in both cases due to their proximity to the cutoff. Figure 9 shows the results of the simulations using the 0.09-0.1 bin. Figure 10 (*n*=40, p=0.113) shows the result of simulations in the first insignificant frequency bin of0.1-0.2.

**Figure 8:**
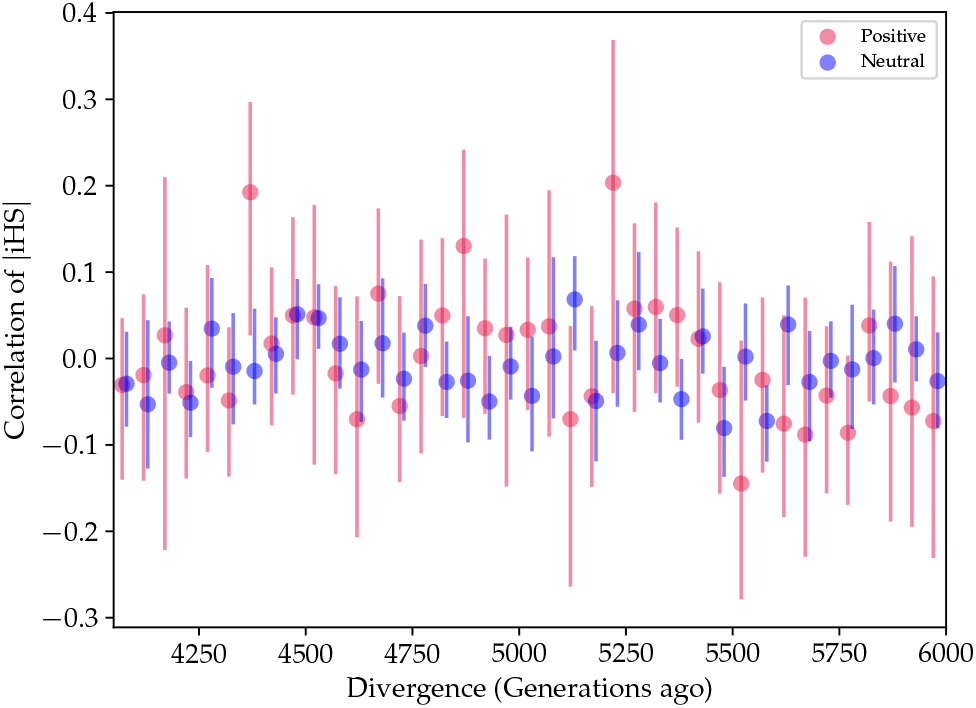
Correlation of |iHS| for a set of divergence times ranging from 4,000 to 6,000 generations ago. No significant difference between neutral and selected regions is detectable.

**Figure 9:**
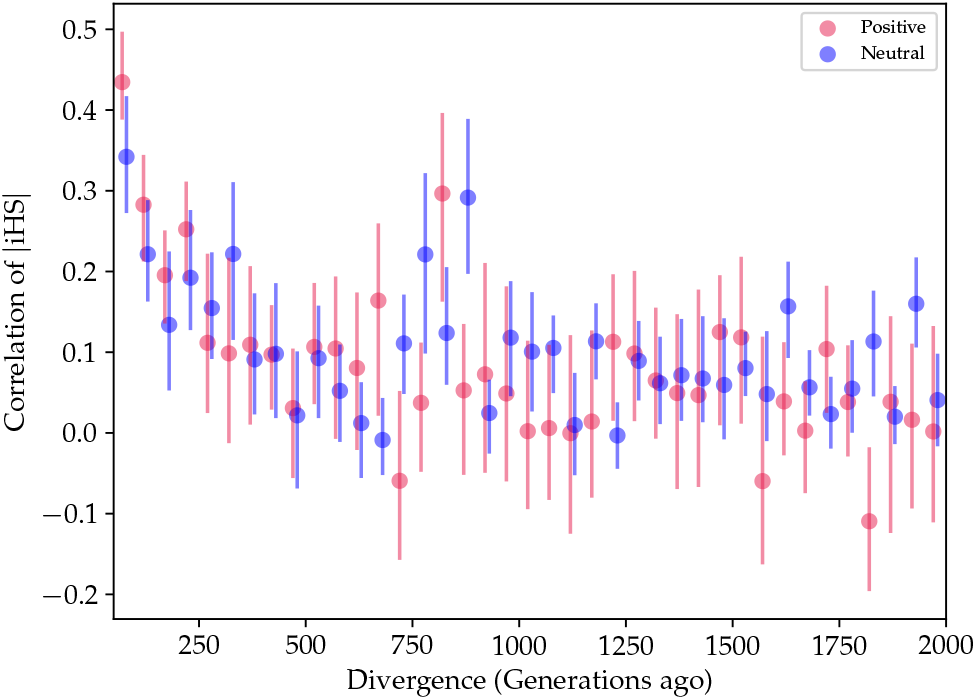
|iHS| correlation with soft-sweeps occurring in one branch with a starting allele frequency between 0.09 and 0.1.

**Figure 10:**
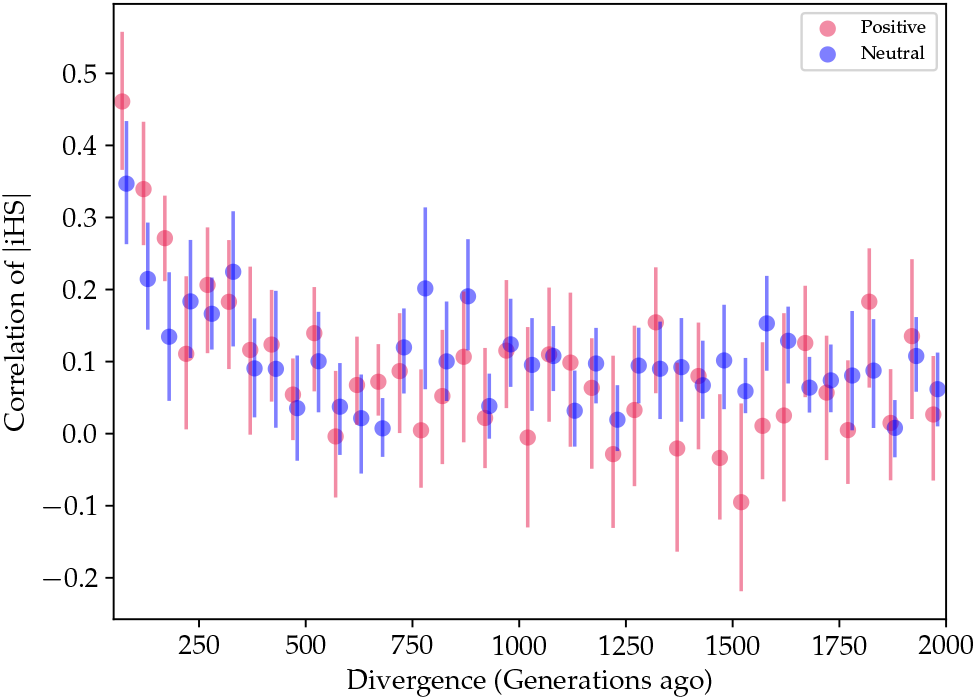
|iHS| correlation with soft-sweeps occurring in one branch with a starting allele frequency between 0.1 and 0.2.

## 3 Discussion

### 3.1 Weak signals differ between populations

The smaller correlation of |iHS| in genic regions compared to nongenic regions implies that populations share few signals of weak selection. This result supports the hypothesis that weak positive selection has a similar disruptive effect as hard classic sweeps. This pattern is also present in the simulated data.

Simulations including beneficial mutations in one population resulted in a depressed correlation of |iHS| compared to neutral simulations. Models of purifying selection did not have the same effect. This suggests that positive selection rather than purifying selection across genic regions is more likely to be driving the increase in population differences.

This effect is amplified when the distance from genic regions increases. If nongenic regions are at least 1Mb from genic regions, confidence intervals for correlation of nongenic regions increase, but the difference between genic correlation and nongenic correlation of |iHS| increases substantially (Figure 11). The sample size as measured by the number of bases and number of regions in the genome both decrease. This change implies two things. First, it supports the notion that nongenic regions are substantially affected by selection at neighboring genic sites. Second, the pattern in the nongenic regions continues to change with increasing distance from genic regions, even to the point that nongenic regions are reduced to a small portion of the genome. This result could be used to support the hypothesis the majority of the genome is affected by positive selection to some extent (e.g., Pouyet et al. (2018); Schrider and Kern (2017)).

**Figure 11:**
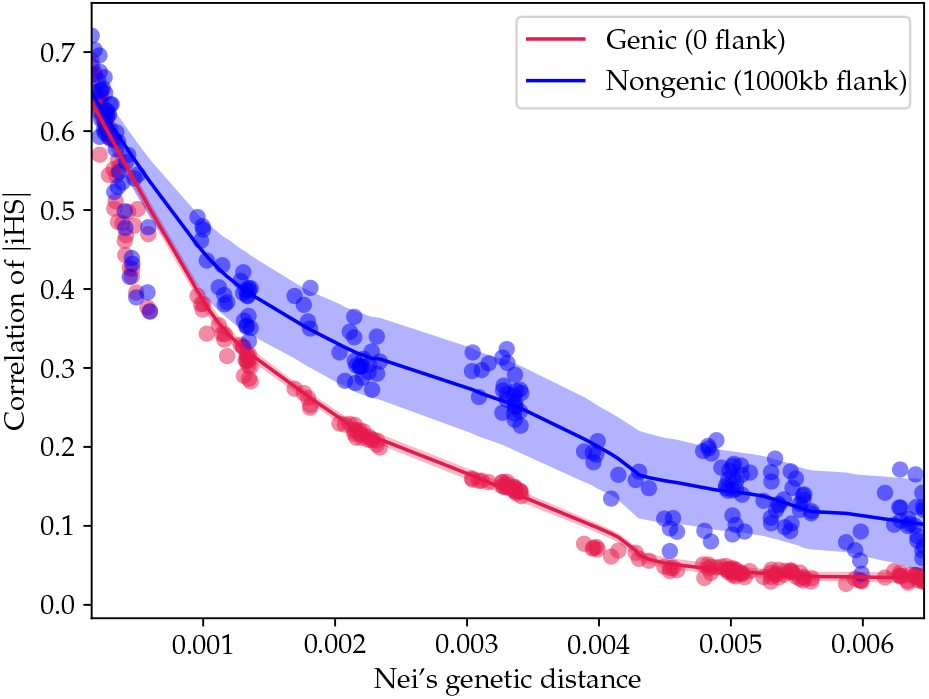
Genic regions compared to nongenic regions with a large flank size of 1Mb.

### 3.2 XP-EHH

The correlation of XP-EHH in genic regions was consistently lower than in non-genic regions, but not significantly. The hypothesis that the patterns in genic regions are due to population history rather than selection cannot be rejected. In other words, the results are consistent with theoretical arguments 1, 2, and 3 enumerated above. However, the XP-EHH results are still informative when compared to |iHS|. The difference in sensitivity between |iHS| and XP-EHH is visible in Figure 12. Pairwise XP-EHH correlation is larger in closely related populations than pairwise |iHS| correlation. This reflects the ability of XP-EHH to detect differences in haplotype structure that are the result of older selection (near fixation) or population history. This implies that the correlation methods implemented here may be repeated for other methods of detecting selection.

Correlations of XP-EHH scores between African populations are substantially small compared to population comparisons with similar genetic distance. This values cluster in the lower left of Figure 12. This is likely due to the increased similarity of these samples to the *reference* XP-EHH sample, Yoruba.

**Figure 12:**
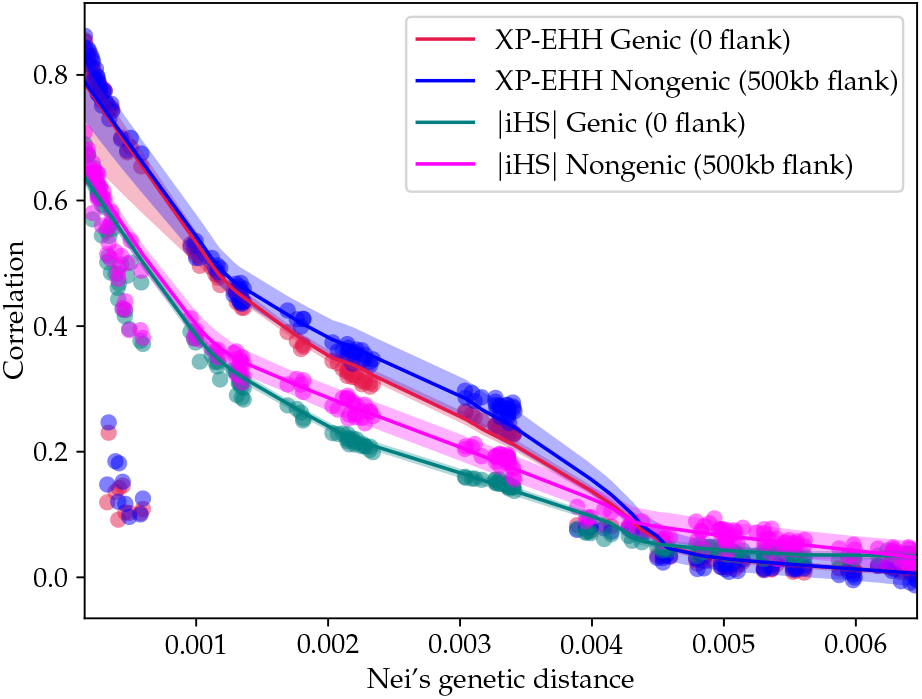
Correlation |iHS| and correlation of XP-EHH scores. Genic regions with population labels can be found in Appendix A.

## 4 Methods

### 4.1 Pairwise correlation of |iHS|

To measure selection, the Integrated Haplotype Score (|iHS|) was calculated for the 1000 Genomes Phase 3 data acquired from ftp://ftp.1000genomes.ebi.ac.uk/vol1/ftp/release/20130502/. Recently admixed populations were excluded from this analysis, leaving 21 samples from Eurasian and African populations. Data were divided by population sample and filtered to exclude sites with a minor allele frequency less than 0.05.

|iHS| was calculated using the *selscan* package (Szpiech and Hernandez, 2014). |iHS| is sensitive to variation in allele frequency (Voight et al., 2006) and local recombination rate (O’Reilly et al., 2008). This was compensated for in two ways. First, when integrated across the chromosome, genetic map distance was used rather than physical distance. Genetic maps for the 1000 genomes were downloaded from the Pickrell lab (https://github.com/joepickrell/1000-genomes-genetic-maps). The map positions for missing sites were imputed from neighboring sites. Second, when the ratio of scores is taken between the two allele types, the effects of recombination ought to disappear because the recombination rate is the same for carriers of both alleles. To verify this process, |iHS| was first standardized using allele frequency bins and then was regressed against recombination rate. A strong relationship between frequency standardized |iHS| and recombination rate remained. Following the example of Johnson and Voight (2018), |iHS| scores were restandardized using 46 frequency and 21 recombination bins. Frequency bins ranged from 0.05 to 0.95 and recombination bins were determined by grouping the data into percentiles.

Standardized |iHS| scores were split into genic and nongenic regions of the genome for each sample using coordinates of known genes and gene predictions from the UCSC table browser (Karolchik et al., 2004). Dividing the data into genic and nongenic regions allows us to distinguish between shared sweeps in a common ancestor or independent sweeps after a common ancestor from spurious signals of selection from common ancestry (theoretical points 1 and 2 from 3 above). Unlike genetic drift, selection affects specific loci and linked variation rather than the entire genome. Genic regions should more commonly be the target of selection because they are more often functional (Barreiro et al., 2008; Coop et al., 2009). The difference between genic and non-genic correlations at a given value of genetic distance can be attributed to selection.

Regions were considered nongenic if they occurred outside of a flanking region around genes. This cutoff is used to compensate for the effects of linkage (Slatkin, 2008; Wall and Pritchard, 2003). To determine the appropriate flank size, correlation of |iHS| was calculated between populations for each subdivision of the genome and increasing flank size. The absolute value of iHS is taken because the sign of iHS only indicates allele state. A beneficial allele will produce both positive and negative scores with large magnitudes at neighboring sites. Correlation was limited to loci with |iHS| values within two standard deviations of zero. This distinction was made to eliminate loci showing potential evidence for strong selective sweeps. The ideal flank size for nongenic regions was assessed based on sample size and the effect of linked genic sites (Table 1). As flanking regions become large, both sample size and the influence of genic regions on nongenic regions decrease. Within genic regions the size of the flanking region had little effect on |iHS| correlations (Figure 13). Therefore, “genic” in this study is defined to mean a flank size of zero. For nongenic regions, a flank size of 500kb was used to balance between sample size and the effect of neighboring genic regions (Figure 14).

**Figure 13:**
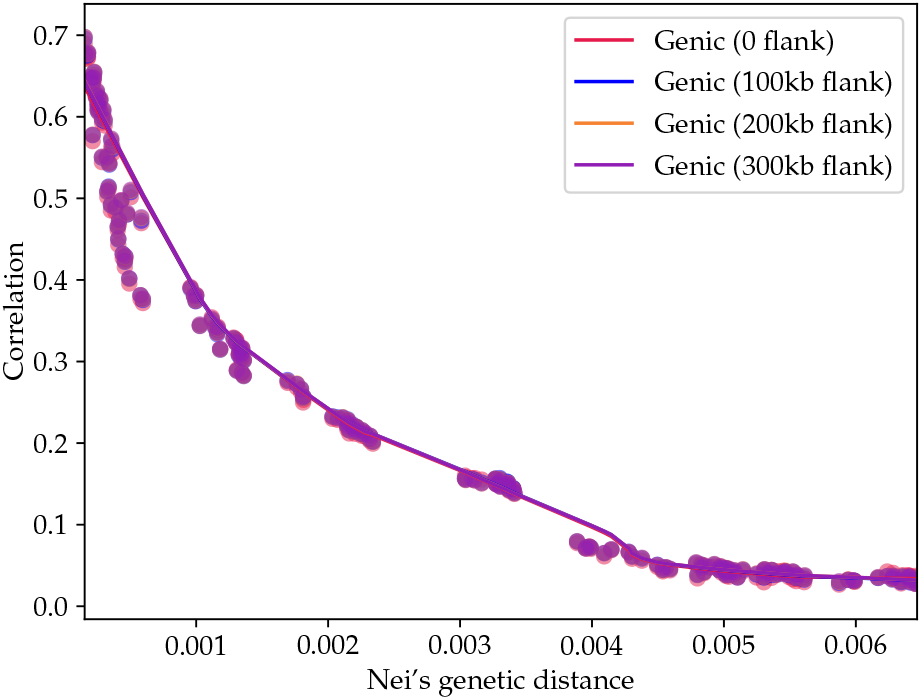
Difference in correlation of |iHS| in genic regions given varying flank sizes.

**Figure 14:**
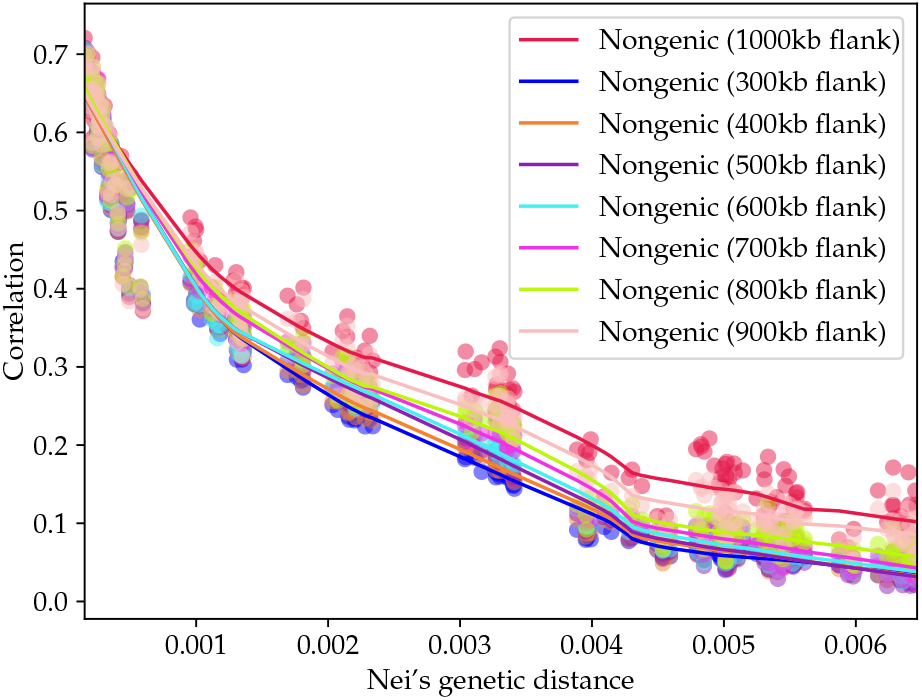
Difference in correlation of |iHS| in nongenic regions given varying flank sizes.

**Table 1:**
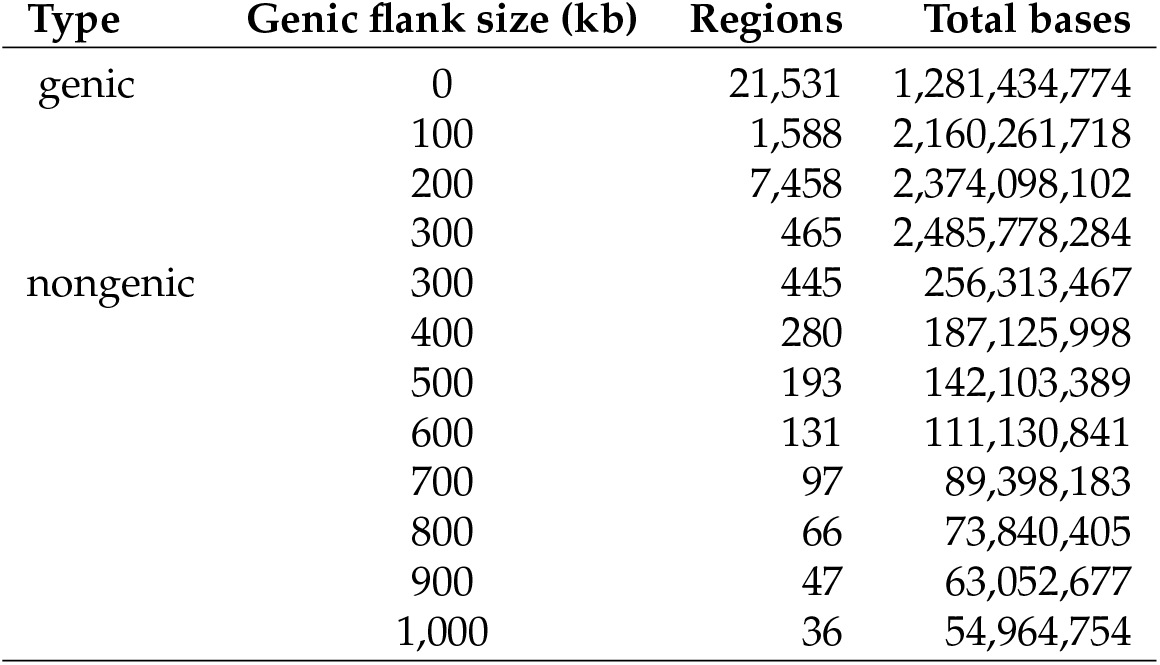
The flank sizes tested for genic and nongenic regions. Genic regions are given flanking regions and nongenic regions are considered to be anywhere not included.

Once flank sizes were determined, genic and nongenic |iHS| correlation matrices of the determined flank size were regressed against Nei’s genetic distance using the Loess algorithm. Nei’s genetic distance (Nei, 1972) was calculated for each pair of population samples using the allele frequencies taken from the 1000 Genomes data.

A moving blocks bootstrap (Liu and Singh, 1992) was used to generate confidence intervals around the regressions. This method was chosen because loci near one another are used in each other’s calculation of |iHS|, implying that |iHS| calculation of neighboring sites is not independent. The moving blocks bootstrap method compensates for this problem by sampling entire regions of the genome rather than individual loci. Blocks of 500 kilobases (kb) were sampled from the standardized |iHS| output. This cutoff was chosen because most blocks of LD in the genome are smaller than 500kb (Slatkin, 2008; Wall and Pritchard, 2003). The number of blocks used in each bootstrap is equal to the number of blocks required to simulate the length of the real data. Correlation of |iHS| was calculated for each resampling. The model fit to the real data is then applied to the resampling of the data. This process is repeated 1,000 times. The inner 95% of these replicates become the confidence intervals for the real data.

### 4.2 Pairwise correlation of XP-EHH

Tests performed on |iHS| were repeated for XP-EHH. XP-EHH requires a second population to serve as a comparison. Results of XP-EHH indicate where selection has occurred in one population or the other, but not both. Each test used the same second or *reference* population, the Yoruba. This will bias the XP-EHH results against signals that populations share with the reference population. However, it allows for the pairwise comparison of nonreference populations. XP-EHH was calculated using the *selscan* package (Szpiech and Hernandez, 2014). XP-EHH scores were standardized using 46 frequency and 21 recombination bins. Frequency bins ranged from 0.05 to 0.95 and recombination bins were determined by grouping the data into percentiles. A moving blocks bootstrap (Liu and Singh, 1992) was used to generate confidence intervals following the same methods described or |iHS| above.

The correlation of XP-EHH was then calculated from the standardized values. Final standardized scores were filtered in both populations to exclude sites that showed evidence for selection in Yoruba. The inclusion of these sites would bias the results. Covariance between populations would be positive in regions where Yoruba experienced selection that was relatively strong compared to the populations being compared. In these regions, both populations would have negative XP-EHH scores with relatively large magnitudes. This increase in covariance would be artificial, rather than reflecting any difference in the populations being compared. The pairwise XP-EHH correlations within Africa clustered at lower correlations than other within continent comparisons. This was visible in Figure 12. Figure 15 shows the same set of results with Africa excluded. The trend in the data does not change in any statistically significant way.

**Figure 15:**
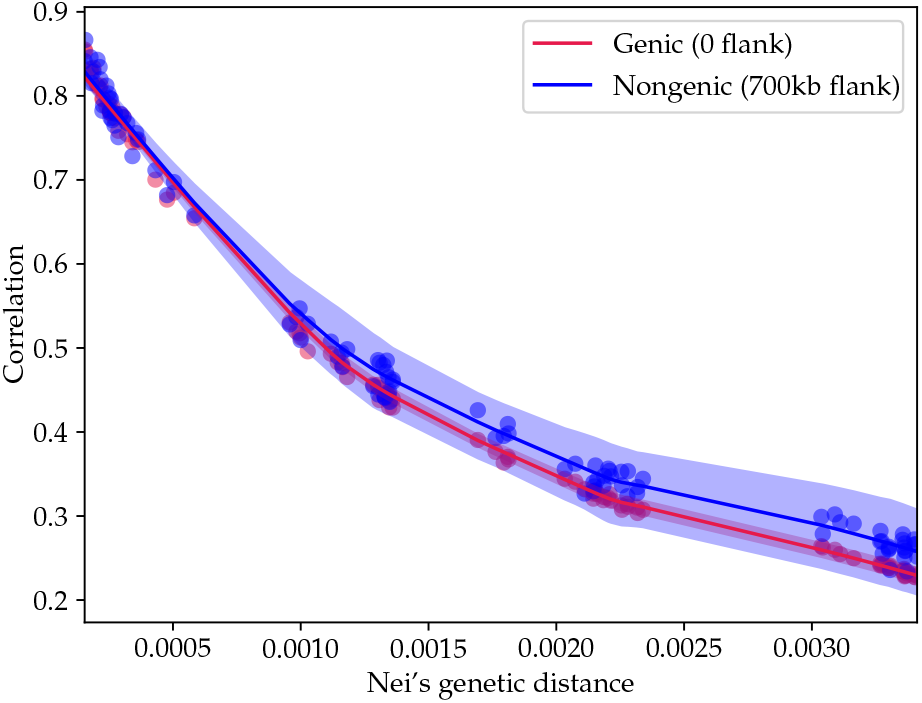
Correlation of XP-EHH scores omitting African samples.

### 4.3 Simulation

Simulated data were generated using SLiM (Haller and Messer, 2018; Messer, 2013). SLiM is a forward-time simulator that allows researchers to model evolutionary scenarios in the presence of linkage. For this work, three basic simulations were performed: a neutral model, a model of purifying selection, and a model of positive selection. Instead of attempting to model human history, a simple model was constructed in which a single population separates into two populations at a prespecified time. This divergence time was adjusted to values between 50 and 2000 generations. A constant population size of 10,000 was used throughout the simulation. The recombination rate was held constant across the simulated genome to eliminate any possibility of an association between linkage disequilibrium and recombination rate. Parameter values can be found in Table 2.

**Table 2:**
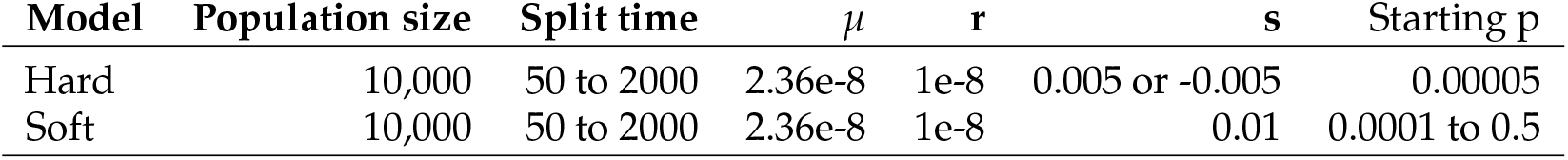
Values used in the simulations

In the neutral case, mutations have no effect. In both cases of selection, selection occurs in one population following a population split. This creates a scenario in which all loci under selection in one population were neutral in the other. Beneficial or deleterious mutations constituted 5% of the total number of mutations in their respective simulations and have a selection coefficient of 0.005 and −0.005, respectively. |iHS| is standardized to eliminate the effects of allele frequency differences caused by drift. In real data, the entire genome is standardized together, and results indicate how exceptional a particular site is given its allele frequency. To replicate this effect in the simulated data, neutral and non-neutral simulations with the same divergence time were standardized jointly for allele frequency.

For each simulation, a moving blocks bootstrap was used to find confidence intervals. A block size of 500 kb was used in the simulations to be consistent with the analysis performed on the real data. Each simulation is independent allowing the use of a sign-rank test. The Wilcoxon signed-rank test was performed between the selection and neutral models.

Simulations of soft sweeps differed from hard sweeps models by using a constant selective coefficient of 0.01, and varied beginning allele frequencies. Soft sweeps start following the population split but occur at manually specified loci meeting the desired allele frequency. This occurs at user-specified in-tervals. In hard sweep models, most mutations, including the beneficial ones, are lost to drift. Soft sweeps starting at higher allele frequencies are unlikely to be lost. This presents a problem because the absolute number of soft sweeps affects the result. Therefore these simulations were run varying the number of introduced of soft sweeps until an allele frequency cutoff was observed.

## Notes

### Competing Interest Statement

The authors have declared no competing interest.

